# Deep-BGCpred: A unified deep learning genome-mining framework for biosynthetic gene cluster prediction

**DOI:** 10.1101/2021.11.15.468547

**Authors:** Ziyi Yang, Benben Liao, Changyu Hsieh, Chao Han, Liang Fang, Shengyu Zhang

## Abstract

Natural products produced by microorganisms constitute an important source of essential pharmaceuticals, including antimicrobial and anti-tumor drugs. These bioactive molecules are microbial secondary metabolites synthesized by co-localized genes termed Biosynthetic Gene Clusters (BGCs). The rapid increase of microbial genomics resources, due to the availability of high-throughput sequencing technologies, has spurred the development of computational methods for microbial genome mining for BGC discovery. Current machine learning methods, however, have limited successes in uncovering novel BGCs due to an excessive number of false positives in their predictions. To this end, we propose Deep-BGCpred, a framework that effectively addresses the aforementioned issue by improving a deep learning model termed DeepBGC. The new model embeds multi-source protein family domains and employs a stacked Bidirectional Long Short-Term Memory model to boost accuracy for BGC identifications. In particular, it integrates two customized strategies, sliding window strategy and dual-model serial screening, to improve the model’s performance stability and reduce the number of false positive in BGC predictions. We compare the proposed model against other well-established methods on common benchmarks and achieve new state-of-the-art results with convincing evidences. We expect that researchers working on genome mining for natural products may be greatly benefited from our newly proposed method, Deep-BGCpred.

## Introduction

Natural products are chemical compounds produced by living organisms. Particularly, bioactive compounds produced by microorganisms constitute a major source of innovative pharmaceuticals including antibacterial, anti-tumor, and immunosuppressive agents (1). To handle challenges posed by the rapidly rising number of clinically drug-resistant bacteria (2, 3), the emergence of new pathogens and viruses (4), and the demand for new drugs for complex diseases (5), researchers have been continuously working to discover novel and structurally diverse bioactive compounds.

Many valuable bioactive compounds are microorganisms’ secondary metabolites, whose synthetic instruction is primarily encoded in a set of genomically co-localized genes known as Biosynthetic Gene Clusters (BGCs) (6–8). While BGC sequences indicate the existence of some potentially bioactive natural products, they still have to be extracted, isolated and purified from microorganisms and experimentally validated (9). With the rapidly evolving high-throughout sequencing technologies, it is progressively easier to collect complete microbial genome sequences. Mining of these genomes has revealed a vast abundance of BGCs, far exceeding the experimentally verified natural products, which indicates that many BGCs are not expressed under laboratory conditions (10). Therefore, the increasing of microbial genomics resources accelerates a data-driven paradigm shift in natural product based drug discovery.

Computational methods play a crucial role in the development of the field of natural product genome mining. Current methods for mining microbial genomes for BGC identification can be roughly divided into two main types: rule-based and machine learning approaches.

The rule-based approach utilizes manually curated “rules” to identify BGCs based on their similarity to reference genes and protein domain composition. The earlier attempts of BGC identification mainly rely on traditional bioinformatics programs, such as BLAST, to align BGC reference with the manual curation (11, 12). For instance, PRISM 4 (13), a comprehensive platform for prediction of BGCs and the secondary metabolite chemical structures from microbial genome sequences, operates in this manner. Another salient example is antiSMASH (14–18), the most widely used rule-based approach for mining microbial genomes for BGCs. Recently, they present antiSMASH version 6.0 (9), which extends the previous versions by expanding and improving the set of BGC detection rules. With the enrichment of the detection rules, it is conductive to improve the accuracy of identifying known BGCs. Although these rule-based approaches are effective for identifying known BGC classes, the known ones account for only a small part of the whole BGC classes, little is known about the vast majority of BGC classes.

The machine learning models offer a new approach to potentially improve the accuracy of BGC detection and discovery in bacterial genome sequences. (19) developed a hidden Markov model-based (HMM) probabilistic algorithm, ClusterFinder, that provides a general solution to the gene cluster identification of both known and unknown classes. Although the HMM-based algorithm is effective, it only remembers partial sequence genetic information and cannot capture high-order interaction patterns between entities (20– 22). This shortcoming severely limits its capability to unambiguously identify all BGCs in thoroughly studied bacterial genome. DeepBGC (23) addresses this limitation by implementing a Bidirectional Long Short-Term Memory (Bi-LSTM) model and protein family (Pfam) (24) domain embedding representations (Pfam2vec) to properly process genome sequence for BGC predictions. Unlike HMMs, DeepBGC is capable of intrinsically capturing short- and long-term dependency between adjacent and distant genomic entities (25). It significantly outperforms ClusterFinder for identifying BGCs in selected samples of bacterial genomes.

Although current machine learning methods provide convincing evidences that it could elevate the state of the art in BGC detection and identification of novel BGC classes in genome sequences (19, 23), few aspects of the current solutions could still be refined in practice. On one hand, they need to tune hyperparameters using a subset of real world data to guarantee that the model trained on the artificial training set performs with the same accuracy on real genomes, which inevitably requires training the model from scratch for a particular application. This tends to raise their application difficulty and compromise their performance stability for real problems. On the other hand, existing methods still rely on a rule-based post-processing to either merge or filter putative clusters in order to suppress the number of false positives in BGC predictions. For instance, DeepBGC introduces a postprocessing step to exclude potentially false-positive regions in the predicted sequence, according to a manually curated rule. This kind of rules, however, could over-simplify the situation and prevent us from uncovering certain BGCs.

In this paper, we propose Deep-BGCpred, a unified framework that effectively addresses the aforementioned customization challenges that arise in natural product genome mining. Deep-BGCpred embeds multi-source protein family domains and employs a stacked Bidirectional Long Short-Term Memory (Bi-LSTM) neural network for an enhanced modeling of various correlations in bacterial genome sequences. Embedding Pfam domains from multiple sources provides additional biological information that complement the Pfam2vec embedding representation nicely. In addition, Deep-BGCpred integrates two customized strategies (detailed in the next section), termed sliding window strategy and dual-model serial screening, to maintain the model’s performance stability in the analysis of the bacterial genome sequence and reduce the number of false positives in BGC identification.

We systematically evaluate the capability of Deep-BGCpred by conducting benchmark experiments on real-word reference bacterial genomes. For the BGC identification, the proposed method outperforms state-of-the-art machine learning approaches by a large margin, which supports the claim that the new model can effectively identify BGCs within genome sequences. Remarkably, the proposed method correctly predicts more BGCs without producing too many false positives that bothers some other machine-learning based methods. We expect that researchers working on genome mining for natural products may be greatly benefited from the newly proposed method, Deep-BGCpred.

## Methods

In this section, we present the proposed deep-learning framework for identifying BGCs from bacterial genomes and classifying them into various natural product classes. The pipeline of genome mining for BGCs is shown in Figure 1, which consists of eight major steps, including input of a genome sequence, gene prediction, protein family domain prediction, prediction of each Pfam domain score, gene-level score summary, candidate BGC identification, prediction of BGC class, and non-BGC filtration. We propose Deep-BGCpred, a deep-learning method for BGC identification within genomes. A main contribution of Deep-BGCpred is derived from our observation that the usage setting for a typical deep-learning model is different between the training and the application stage (i.e. testing stage) in this field. Specifically, the number of Pfam domains in a typical data instance is inconsistent between the artificially created training set and the real genomic data. Hence, we propose the sliding window strategy to bridge the differences in the usage setting under these two scenarios. Another contribution is the adoption of a dual-model serial screening that could suppress the number of false positive in BGC predictions. In the rest of this section, we elaborate on details of the 8-step workflow summarized in Figure 1.

**Fig. 1.**
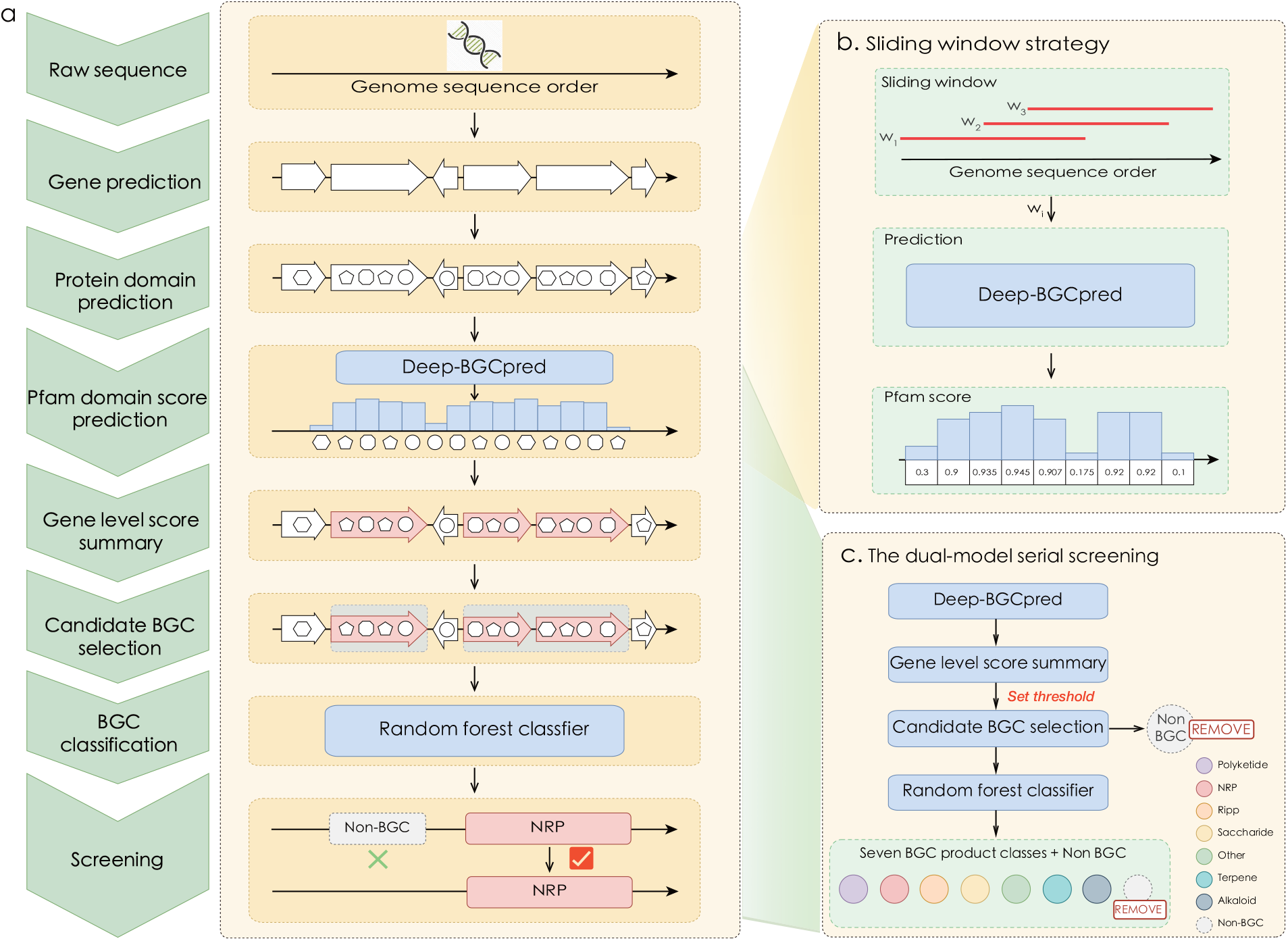
(a) A novel framework for identification of Biosynthetic Gene Clusters in bacterial genomes. (from top to bottom) raw genomic sequences (solid line) are used for gene (arrowhead structures) prediction using Prodigal (26). Pfam domains (circles, Pentagons, and hexagons) are annotated to each genes/open reading frames using hmmscan (27). Deep-BGCpred predicts the classification score for each Pfam domain (blue bar). Pfam domain scores are summarized across genes, which are selected accordingly (red arrowheaded structures). Consecutive candidate BGC genes are assembled into the putative BGCs (blue rectangles). BGCs are classified using random forest classifier based on compound class. Non-BGC regions (gray dashlined rectangles) are filtered out by the screening strategy. (b) Sliding window strategy. Multiple Pfam sequence fragments are extracted from the raw genome sequence by a sliding window as the input of Deep-BGCpred. (c) Dual-model serial screening. Utilizing two methods, Deep-BGCpred and random forest classifier, to jointly reduce the false positive detections in a serial way.

### Gene prediction with annotation of open reading frame

Open reading frames (ORF) of the raw genomic sequence are predicted by using Prodigal (26) version 2.6.1 with the default parameters.

### Protein family identification

Protein family domains are identified using hmmscan (27) version 3.1b2 against the Pfam (protein family) database version 31 (24). The hmmscan tabular output is filtered using BioPython SearchIO module (28) version 1.70, keeping only the highest scoring Pfam regions with evalue *<* 0.01. The resulting list of the Pfam domain is sorted according to the gene and the starting position of the domain.

### Deep-BGCpred training set

The positive training set for Deep-BGCpred is curated from the Minimum Information about a Biosynthetic Gene cluster (MIBiG) database version 1.5 (29), which contains 1,984 BGC sequences from 1,094 bacterial species. The negative samples of the training set are generated according to the protocol outlined in (23) and (19). In this case, we generate 10,128 non-BGC sequences as negative samples. The detailed information of BGC training set is provided in Table S3 in the supplementary material. There are 96,412 Pfam domains in the positive samples (i.e. BGC sequence) such that each BGC contains an average of 48.59 Pfam domains, indicating that most BGC sequences have only a few Pfam domains (see Figure S1 in the supplementary material). A total of 9,633 unique Pfam domains are annotated in the training set. The Pfam identifier (e.g., PF00008), Pfam summary information (e.g., PF00008: EGF-like domain) and Clan identifiers (e.g., PF00008: CL0001) for each protein family domain are also recorded in the training set.

### Deep-BGCpred implementation

The proposed Deep-BGCpred method is inspired by the work of DeepBGC (23), which translates a genome sequence into pfam2vec embedding vectors that are subsequently fed into a Bi-LSTM neural network for prediction. In addition to the Pfam2vec vectors, the proposed method also transforms the Pfam domain summary information and the Clan identifiers into the continuous vector representations. All these embedding vectors are concatenated before feeding into the stacked Bi-LSTM neural network, which is responsible to transform these input features into predicted Pfam domain scores.

Deep-BGCpred is implemented using Keras version 2.1.6 with TensorFlow backend version 1.15.4. The overall architecture of Deep-BGCpred is shown in Figure 2. It consists of two parts: Pfam domain encoder and the stacked Bi-LSTM model. Each Pfam domain is first transformed into an embedding representation in the real-valued vector form. Three types of Pfam domain information are encoded. First, the Pfam domain identifiers are embedded into 102-dimensional vectors via pfam2vec released by (23). Second, the Pfam domain summary information are embedded into 960-dimensional vectors via a character-level convolutional neural network (30). Third, the Clan identifiers are embedded into 64-dimensional vectors through a Bi-LSTM neural network. For detailed information about building the Pfam domain encoder, please see the supplementary material. For the stacked Bi-LSTM model, it contains three sub-layers, namely the stacked Bi-LSTM layer, the pooling layer, and the dense layer. The stacked Bi-LSTM layer consists of a stateful Bi-LSTM layer with 128 hidden units and dropout of 0.2 followed by a forward LSTM layer with 128 hidden units. The average pooling layer combines the local feature vector with the previous layer to obtain the global feature vector. The dense layer is composed of a time-distributed dense layers with a sigmoid activation and one output unit.

**Fig. 2.**
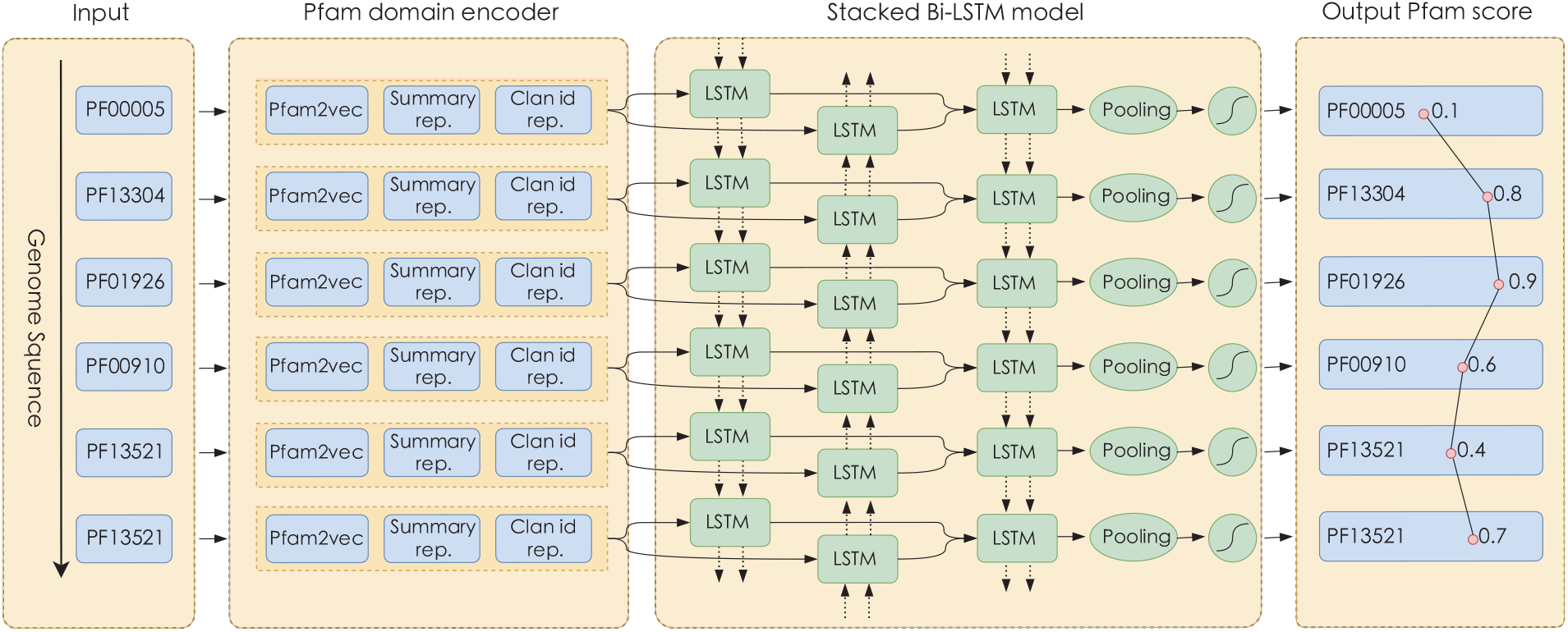
The overall architecture of the proposed Deep-BGCpred method (left to right blocks). It consists of four parts: input, Pfam domain encoder, the stacked Bi-LSTM model, and output layer. Each row represents a timestep in which Deep-BGCpred processes a single Pfam domain in the input sequence, which is maintained in genomic order. Three types of Pfam domain information are embedded into the vector representation: Pfam domain identifier, Pfam domain summary information, and Clan identifier. These embedding vectors are concatenated and fed into the stacked Bi-LSTM model. In each timestep, output from the stacked Bi-LSTM model is processed through a fully-connected node with sigmoid activation function that outputs the score of a given Pfam domain.

The output of Deep-BGCpred is a sequence of values, normalized between 0 and 1, corresponding to the predicted scores for a sequence of Pfam domains to be part of a BGC. In each training epoch, all positive and negative samples are shuffled randomly and concatenated to create artificial genomes, where BGC is randomly scattered throughout the genome and surrounded by non-BGC sequences. Training is configured with 256 timesteps and a batch size of 64. Thus, the training sequence of each epoch is separated into 64 subsequences, each trained in parallel in batches of 256 timesteps, processing a single training vector at each time step. The final model is trained for 100 epochs and optimized via Adam optimizer using a fixed learning rate of 10−4 and weighted binary cross-entropy loss function.

To obtain the BGC region, the predicted score are first averaged in each gene, then BGC genes are selected according to a given threshold, and finally the consecutive BGC genes are combined. Furthermore, the BGC region is filtered by applying the post-processing criteria defined in (19): (1) merge BGC regions separated by at most one gene; (2) filter out the BGC region with less than 2000 nucleotides; (3) eliminate 133 regions with no known biosynthetic domains published in the ClusterFinder (19) submodules of antiSMASH (9).

### Random forest training set

The putative BGCs predicted by Deep-BGCpred are then classified using the random forest classifier. Different from the work in (23), eight classes are classified by the random forest classifier: non-BGC class, and seven classes of BGC based on biosynthetic product (i.e. Alkaloids, non-ribosomal peptide (NRP), Polyketide (PK), ribosomally synthesized and post-translationally modified peptides (RiPP), Saccharide, Terpene, and other unclassified BGCs).

To further reduce the number of false positives in predictions, we build a customized dataset for training the random forest classifier. The biosynthetic product classes are extracted from the MIBiG database version 1.5, producing 2,018 labelled training samples. The non-BGC samples are taken from two sources: data generation based on the negative samples released by (23), and the non-BGC samples incorrectly predicted by Bi-LSTM network in the preceding stage of the training process. To generate negative samples (from the first source), we adopt a novel data augmentation technology, using the Pfam domain similarity network in the EMBL database (31) to extract negative samples. Specifically, the Pfam domains in the non-BGC sequence are randomly replaced by other similar Pfam domains, which is analogous to the synonym replacement in natural language processing (NLP) (32). The probability of a Pfam domain in the sequence being replaced is equal to *max*(2/sequence length, 0.02). In other words, we force at least two Pfam domains to be replaced in the sequence. 2,000 negative samples are generated using this simple yet universal technique. In the end, a total of 2,102 non-BGC samples are collected for training the random forest classifier. Tables S4, S6, and S7 summarize a brief description of the random forest training set.

### Sliding window strategy

The Bi-LSTM network is trained with 1,984 positive and 10,128 negative samples described above. Since this training set is artificially created, the negative samples bear certain characteristics distinct from that of the real-world data (e.g. real distribution and number of the annotated Pfam domains) (23). Current machine learning methods need to tune hyperparameters using a subset of realworld data to guarantee that the model trained on the artificial training set would perform with a similar accuracy on real genomes, which inevitably requires training the model from scratch for a particular application. This, however, tends to raise their application difficulty.

We focus on alleviating the inconsistency of the number of Pfam domains between the artificial created training set and real-word data. When analyzing real genomes, tens of thousands of Pfam domains are typically fed into the model simultaneously to predict whether it is part of the BGC. However, each sample in the artificially created training set contains only 256 Pfam domains. Obviously, this is an inconsistency in the typical traits of training data and real-world data. From the machine learning perspective, this inconsistency may compromise a model’s performance stability in real-world applications.

Sliding window strategy aims at intercepting multiple Pfam sequence fragments from continuous Pfam sequences to ensure a consistency of how we process data during training and real-world applications, without requiring excessive adjustment of hyperparameters in the real-world application. The flowchart of predicting the Pfam score based on the sliding window strategy is shown in Figure 3. First, multiple Pfam sequence fragments are extracted from a continuous Pfam sequence using a sliding window, where the window size is 256 and the step size is 10. Second, each Pfam sequence fragments are then fed into the Bi-LSTM network to obtain the BGC classification score for each Pfam domain. Third, the final score of Pfam domain is obtained by calculating the average value of Pfam domain at each site in genomic order.

**Fig. 3.**
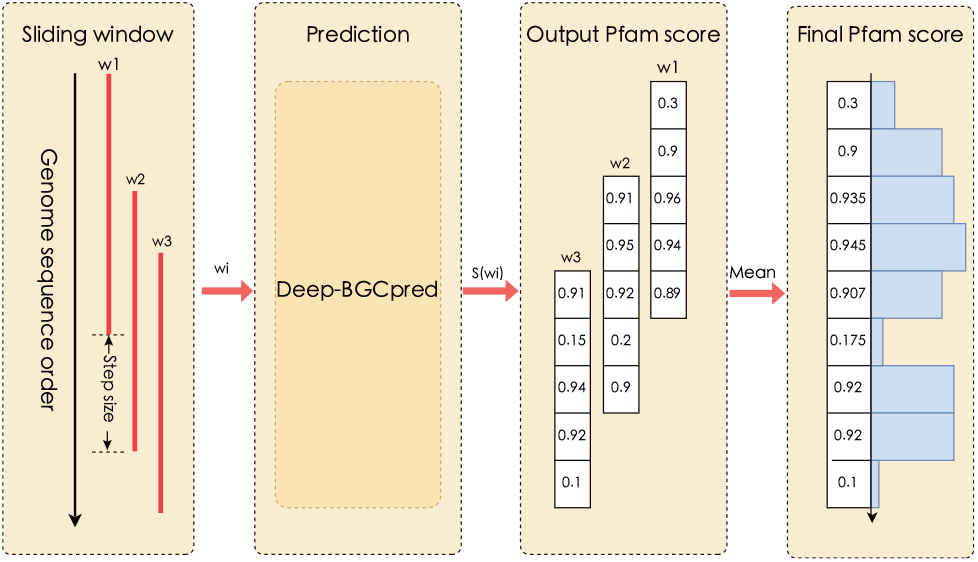
The flow chart of Pfam score prediction based on the sliding window strategy.

### The dual-model serial screening

In addition to excluding BGC regions with no known biosynthetic domains, the proposed Deep-BGCpred framework performs dual-model serial screening to reduce the number of false positives. As shown in Figure 1(c), we adopt the proposed Deep-BGCpred method and random forest classifier to jointly reduce the false positive rate in a serial way. Specifically, this strategy can be divided into two steps. The first step is to use Deep-BGCpred to output the prediction score of a given Pfam domain, summarize the Pfam score at the gene level, and then select BGC genes with any given threshold and merge consecutive BGC genes. The second step is to predict the class of the BGC regions using the random forest multi-label classifier. In both steps, the samples predicted as non-BGC are filtered. Even the samples predicted as BGC in the first step are likely to be removed by the random forest classifier in the second step. The random forest classifier specially learns the effective information of non-BGC samples that Deep-BGCpred has not learned. Specifically, the negative samples predicted incorrectly in Deep-BGCpred are fed into the training pool of the random forest classifier for learning. On the other hand, the first step can only predict whether a sequence belongs to BGC, while the second step can classify the BGC in more detail (i.e. Non-BGC class or one of the seven biosynthetic products), which illustrate this strategy achieves the BGC classification from coarse-grained to fine-grained.

## Results and Discussion

### Deep-BGCpred validation on the BGC sequence dataset

Deep-BGCpred is built upon a strong-performing DeepBGC (23) to further improve BGC identification. To ensure adequate comparison between DeepBGC and our proposed deep learning method, we train and validate Deep-BGCpred with the BGC sequence dataset constructed by (23). The BGC sequence dataset is randomly divided into two sets: approximately 95% of the samples are used for training and the remaining ones are for test.

Figure 4 plots the tendency curves of the training performance calculated on the BGC sequence dataset, as well as those calculated simultaneously on test data during learning iterations. From the figure, we can clearly find that the test accuracy and recall of Deep-BGCpred evidently outperforms DeepBGC after 10 epochs. In particular, the proposed method can obtain a relatively high test recall after 10 epochs, and attains more than 20% gain compared with DeepBGC. It implies that the proposed method identifies more BGCs.

**Fig. 4.**
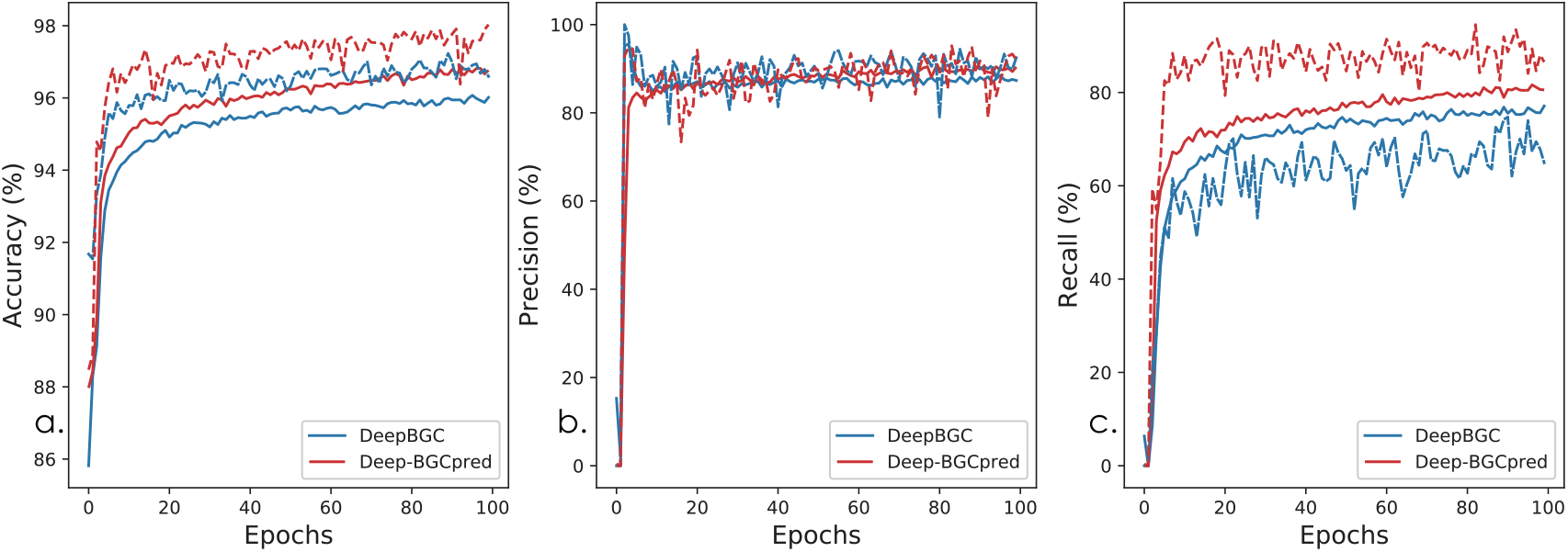
Training and test performance changing curves on the BGC sequence dataset. Solid and dotted curves denote the training and test performance, respectively. (a) Accuracy. (b) Precision. (c) Recall.

### Deep-BGCpred validation on real bacterial genomes

To illustrate the capability of the proposed framework, we curate twelve whole bacterial genomes containing 256 manually annotated BGC regions from (19) for BGC identification. Raw annotated results for each Pfam domain are parsed from the Genebank output file. The detailed information of annotated BGC regions in twelve bacterial genomes is shown in Table S8 in the supplementary material. The comparison methods include: DeepBGC (23); ClusterFinder_original (19), the original ClusterFinder HMM model; ClusterFinder_retrained (23), ClusterFinder HMM retrained with up-to-date data. Deep-BGCpred integrates sliding window strategy and dual-model serial screening for BGC identification.

We evaluate the performance of the proposed framework and compare it with the baseline methods by testing its ability to (1) accurately identify BGC position (i.e. Pfam domain level) throughout the whole bacterial genomics, and (2) distinguish BGC from non-BGC sequence (i.e. BGC level). First, to evaluate the capability of the machine learning methods to accurately identify BGC positions within the whole bacterial genomes, we conduct experiments on Pfam domain level. Figure 5 (a) plots the Pfam domain level Receiver Operating Characteristic (ROC) curve calculated on 12 manually annotated reference bacterial genomes. We can clearly find that the proposed framework achieves the best performance. It achieves AUC with 0.942, which is superior to the results of Clusterfinder_original and Clusterfinder_retrained with 0.839 and 0.916, respectively, and outperforms Deep-BGC by a small margin. However, most of the annotated Pfam domains in the real bacterial genome do not belong to BGC, which is highly imbalanced. Precision Recall Curve (PRC) is more informative than the ROC curve when evaluating the models on imbalanced datasets. The Pfam domain level PRC calculated on 12 annotated genomes is shown in Figure 5 (b). As shown, Deep-BGCpred outperforms all the competing methods by a large margin at Pfam domain level. The average precision (AP) of Deep-BGCpred holds roughly a 5% lead over the second-best result. It implies that Deep-BGCpred predictions are of high precision and are composed of less false positives at Pfam domain level. Moreover, we report the Pfam domain level performance for each competing methods under a certain threshold. Table **??** shows the Pfam domain level performance comparison between all competing methods under threshold= 0.9 setting. It can be observed that our method gets the best performance across almost all evaluation metrics, except the second for recall. Although recall of our method slightly worse than Clusterfinder_original, Deep-BGCpred outperforms Clusterfinder_original on precision by a large margin. Deep-BGCpred attains more than 35% gain compared with Clusterfinder_original. This implies that many ClusterFinder predictions are false positives.

**Fig. 5.**
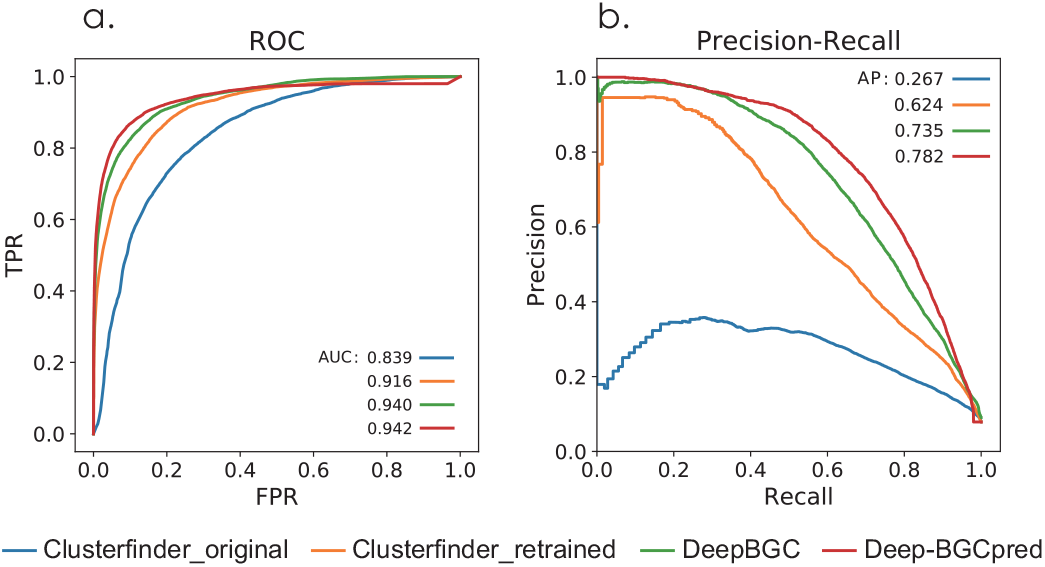
All competing methods validation on the Pfam domain level using the (a) Receiver Operating Characteristic (ROC) curves and (b) Precision Recall Curve (PRC). For ROC curves, the False Positive Rate (FPR) on the *x*-axis and the True Positive Rate (TPR) on the *y*-axis.

Second, we evaluate whether the proposed framework could accurately discriminate between BGC and non-BGC sequences compared with all competing methods. True BGC coverage of each method is calculated for each annotated true BGC region as the fraction of the region that is covered by all its overlapping predicted BGC regions of a given method. A BGC region is considered a true positive when its coverage rate is above a given coverage threshold. Given a coverage threshold, each predicted BGC region is considered as the true positive if it overlaps with a true BGC region and as a false positive if it does not. We adopt the BGC-level F1 score as the evaluation metric. BGC-level precision is calculated by dividing the number of true positives by the total number of predicted BGCs, and BGC-level recall is calculated by dividing the number of true positives by the total number of true BGCs. The BGC-level F1 score is calculated based on BGC-level precision and BGC-level recall. Figure 6 shows the BGC-level F1 score comparison between all competing methods under fixed coverage rate setting. As shown, our method achieves the best performance across all coverage rates. The superiority of Deep-BGCpred is evident. The performance gaps between our method and DeepBGC increase as the coverage rate is decreased from 60% to 1%. Besides, our method outperforms ClusferFinder_orginal and ClusterFinder_retrained by a large margin in all coverage rate tested. It implies that our method can accurately distinguish BGC region and non-BGC region compared with the competing methods.

**Fig. 6.**
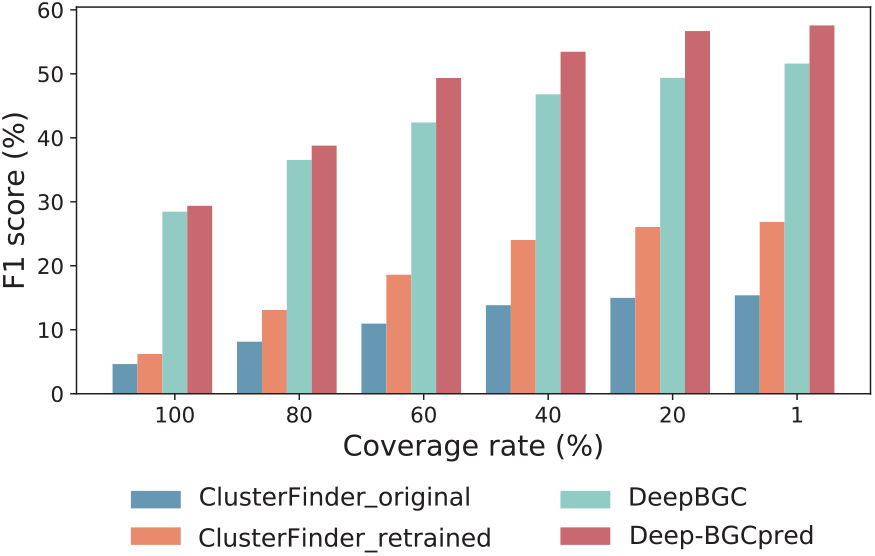
BGC-level F1 score comparison on real annotated bacterial genome dataset with varying coverage rate settings.

**Table 1.**
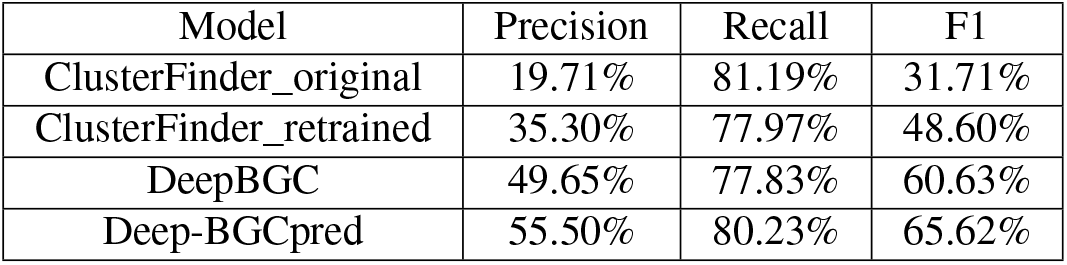
Pfam domain level performance comparison on competing machine learning methods under threshold= 0.9.

### Deep-BGCpred compared with the rule-based methods

To further illustrate the capability of the proposed framework, we compare Deep-BGCpred with the rule-based methods on the twelve manually annotated genome dataset. The compared methods include: antiSMASH version 6.0 (9) and PRISM 4 (13). It is crucial for us to accurately identify true positive regions, such that we adopt the BGC-level recall as the evaluation metric to compare the ability of all competing methods in identifying true BGC regions.

Figure 7 shows the BGC-level recall of the two types of methods on real bacterial genome dataset with varying coverage rate. The machine learning method to identify BGC is threshold-dependent, selecting BGC genes by a given threshold and then identifying potential BGC regions. Here, we present the results of machine learning methods with the threshold = 0.7, 0.8, 0.9, respectively. Since most of the ClusterFinder predictions are false positives, we do not present its predicted results in this subsection. As shown in Figure 7, with the decrease of threshold, the BGC-level recall of Deep-BGCpred and DeepBGC are improved. For instance, when coverage rate= 60%, the BGC-level recall of Deep-BGCpred is 65.63%, 72.66%, and 76.95% with thresholds 0.9, 0.8, and 0.7, respectively. When threshold= 0.7, the proposed method outperforms all competing methods by a large margin on all coverage rate settings. The results suggest that the advantage of the machine learning method is that the threshold can be adjusted to enhance the overall accuracy of uncovering novel BGC regions. In addition, we can observe that the BGC-level recall of the machine learning methods rise more quickly than that of the rule-based methods as coverage rate decreases. The superiority of Deep-BGCpred is evident when the coverage rate is lesser than 60%. It implies that Deep-BGCpred excels at identifying novel BGC regions compared with all competing methods.

**Fig. 7.**
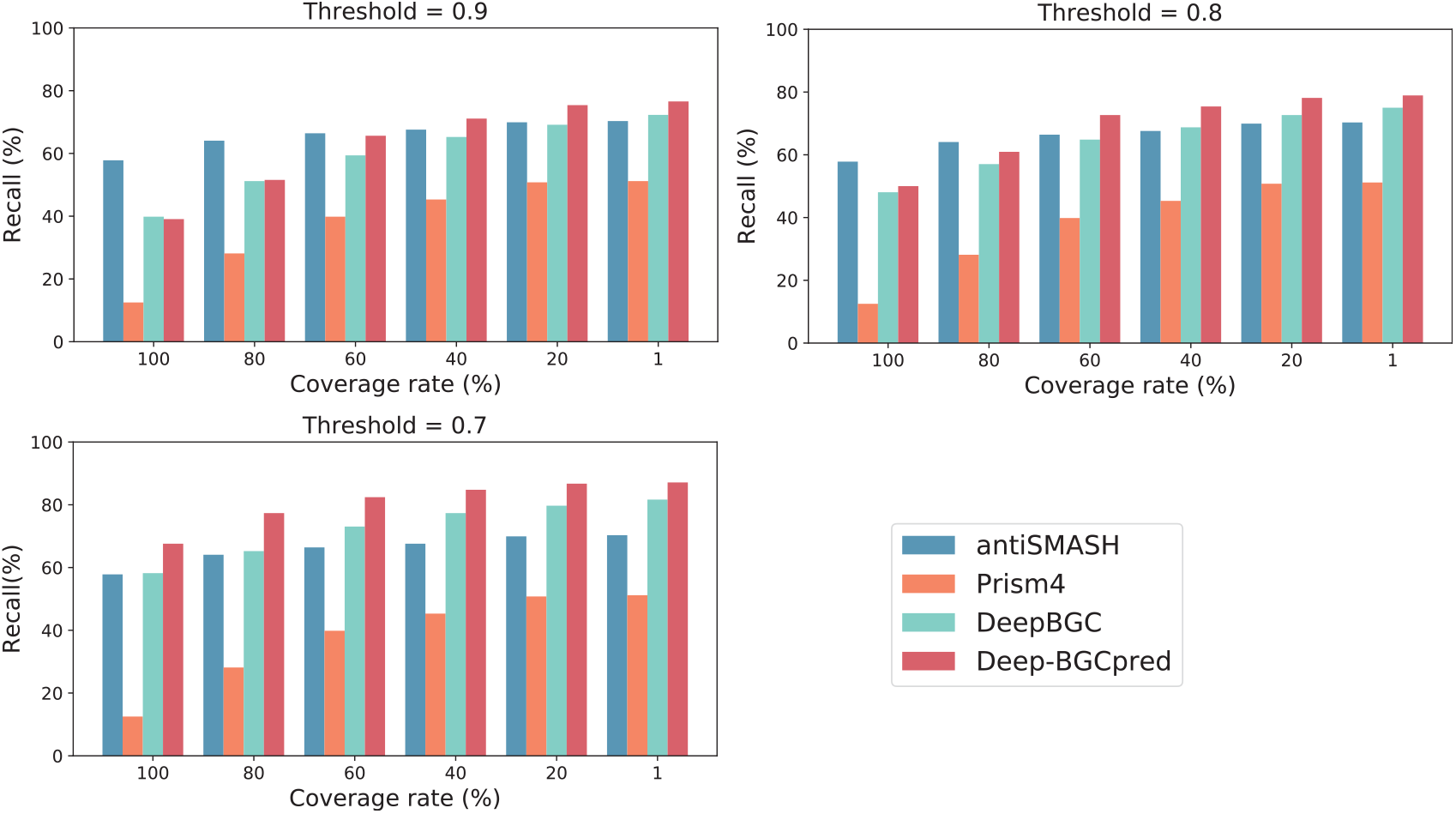
BGC-level recall comparison on real annotated bacterial genome dataset with varying coverage rate settings. Two types of method: rule-based and machine learning method. The machine learning method is threshold-dependent. The threshold settings are 0.7, 0.8, and 0.9, respectively.

We plot the whole genome view of the true annotated BGC regions and the predicted BGC regions by all competing methods on twelve annotated bacterial genomes, as shown in Figure S2 in the supplementary material. For simplicity, only part of the contig is displayed in Figure 8. As shown in the figure, PRISM 4 identifies the least number of BGCs and has the lowest false positive rate. This result also indicates that PRISM 4 has a relatively weaker ability to identify novel BGC regions. Compared with PRISM 4, anti-SMASH exhibits better capability to identify the true positive regions. However, it cannot detect the novel BGC region when a certain BGC region does not conform to the human-coded rule sets. From the snapshot of Streptomyces ghanaensis, we observe that there are two BGC regions identified by Deep-BGCpred but missed by all other competing methods. It implies that our method has the potential for identifying previously unknown sources for natural products in existing bacterial genome sequences.

**Fig. 8.**
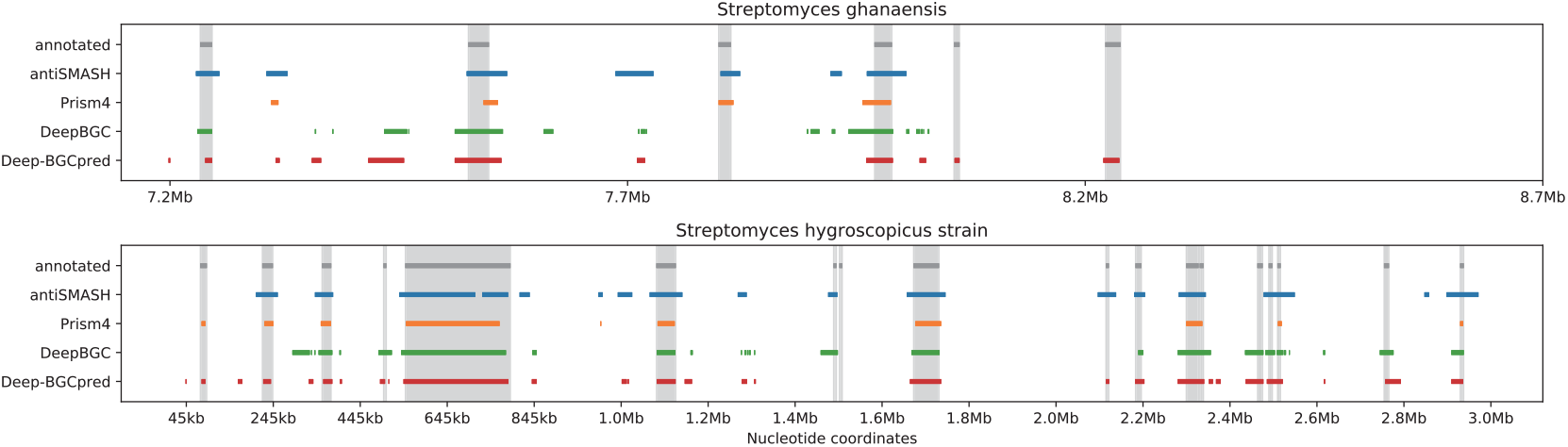
A snapshot of the partial contig view on two manually annotated bacterial genomes. x-axis indicates the genomic coordinates of the bacterial, manually annotated BGCs (grey shade and bar), antiSMASH (blue bar), Prism4 (yellow bar), DeepBGC (green bar), Deep-BGCpred (red bar). The threshold is set as 0.9 for the machine learning methods.

## Conclusion

In this paper, we introduce the unified Deep-BGCpred framework for BGC identification from bacterial genome sequences. Deep-BGCpred is built upon a strong-performing DeepBGC to further improve BGC identification. In addition, two customized strategies, sliding window strategy and dual-model serial screening, are integrated into the proposed framework to boost the model’s performance stability and reduce the number of false positives in predictions. We apply the proposed framework to the real-world reference bacterial dataset to verify is ability to predict BGC genomic positions and identify BGC regions. Our empirical results show that the proposed method outperforms other commonly used machine learning methods, DeepBGC and ClusterFinder. In particular, We demonstrate that the proposed method can identify new BGCs that are missed by all other existing approaches (rule-based and machine learning methods). This is important as it implies that the proposed method can uncover previously unknown sources of natural products in the existing bacterial genome sequences. By incorporating the two customized strategies, we fix a common problem that existing ML-based solutions tend to wrongly recognize a long BGC as few fragmented and shorter BGCs. In summary, the proposed method establishes a new state of the art on the common benchmark in the field of BGC mining. Particularly, the method manifests a strong potential to be capable of uncovering unknown BGCs, which would offer valuable insights for the search of new natural products.

## Supplementary Note 1: Supporting information Available

The Supplementary Material is available on the “Supplementary Material.docx” file.

**Data S1:** Detailed information on the data set, provided as an Excel spreadsheet (XLSX).

